# G2T: Tissue Reconstruction from Gene Expression via Embedding-Distance Flow Matching

**DOI:** 10.64898/2026.08.25.746917

**Authors:** Sebastian Birk, Fabian J. Theis, Mohammad Lotfollahi

**Author notes:** Joint last authors (Fabian J. Theis, Mohammad Lotfollahi).

## Abstract

Single-cell RNA sequencing (scRNA-seq) profiles transcriptomes at high resolution but discards the spatial context of cells within a tissue—information that is essential for studying intercellular mechanisms and tissue architecture. Spatial transcriptomics (ST) retains coordinates but, depending on the assay, trades this off against gene-panel breadth, spatial resolution, or cost. We present **G2T** (Gene-to-Tissue), a generative deep learning model that reassembles a tissue from gene expression — its only observed input — by predicting the matrix of pairwise distances between cells in a learned embedding space. G2T uses an attention-based Transformer with an Euclidean-Distance-Matrix (EDM) output head and is trained with conditional flow matching: the network learns to denoise corrupted cell positions, conditioned on the slice’s gene expression, by predicting per-cell embeddings whose pairwise squared distances match the ground-truth distance matrix. At inference, a fast locally-optimal-block (LOBPCG) multidimensional scaling step turns the predicted distance matrix into 2-D coordinates. On a published MERFISH mouse primary motor cortex benchmark, G2T improves over the previous state-of-the-art method, LUNA, across all three standard metrics— Spearman correlation of pairwise-distance ranks, Contact F1, and per-cell-class Sum RSSD — and even larger relative gains on the mouse central-nervous-system scRNA-seq atlas, evaluated against an imputed spatial reference (STARmap PLUS-integrated locations, not measured coordinates). By predicting this geometry in a higher-dimensional embedding space rather than regressing 2-D coordinates, G2T relaxes the 2-D output parameterisation of prior diffusion-based methods and yields a compact, scalable building block for reconstructing tissue from dissociated cells, enabling downstream spatial niche and cell–cell communication analysis.

## 1. Introduction

Single-cell RNA sequencing (scRNA-seq) profiles transcriptomes at cellular resolution, but dissociation discards each cell’s spatial context. Spatial transcriptomics (ST) retains coordinates, but current assays trade off against one another: imaging-based methods (e.g. MERFISH) profile single cells across only a targeted gene panel, whereas sequencing-based methods are transcriptome-wide but trade off spatial resolution or per-cell sensitivity — and both remain lower-throughput and costlier than dissociated scRNA-seq. Most large atlases produced today are therefore dissociated, and the spatial information that downstream analyses such as niche identification [19], cell–cell communication [4], and tissue-architecture analysis [13] depend on must be re-introduced computationally.

Existing computational methods broadly fall into two families. *Reference-mapping* methods (Tangram [1], CytoSPACE [18], STEM [3], scSpace [14], STALocator [9], CeLEry [27]) map dissociated cells against a fixed ST reference slice, and their accuracy depends strongly on which slice is chosen [24]. *Reference-free* methods reconstruct geometry without committing to a single slice: novoSpaRc [12] casts reconstruction as optimal transport under the assumption that physically proximal cells tend to share similar expression, and D-CE [28] embeds a cell–cell transcriptomic network to recover spatial organisation, while CellContrast [10] and COME [22] learn contrastive embeddings in which embedding distance reflects physical proximity, and a distance-preserving VAE [29] reconstructs coordinates by regularising latent distances toward spatial ones. Most prominently, LUNA [24] frames reconstruction as conditional generation, training a diffusion model with an SE(2)-invariant pairwise squared-distance loss to reassemble tissues of more than a million cells.

LUNA establishes pairwise-distance supervision as a powerful objective for tissue reassembly, because rigid motions of a slice are all biologically equivalent. We inherit this recipe — the SE(2)-invariant pairwise squared-distance objective, the Transformer denoiser, and conditioning on the generative time and the noisy positions — and change *what* the denoiser predicts: rather than regress 2-D coordinates directly, we reparameterise it with an Euclidean-distance-matrix (EDM) head that emits per-cell embeddings in ℝ^*K*^ whose pairwise squared distances form the predicted geometry, recovering 2-D coordinates by a fast multidimensional-scaling (MDS) step. Training uses conditional flow matching [11] (straight-line probability paths, no hand-designed noise schedule); in our ablations it matches the diffusion objective on accuracy while cutting sampling from 1,000 to 50 steps (Table 1), so we treat it as a sampling-efficiency choice rather than the source of the accuracy gain. The resulting model, **G2T** (Gene-to-Tissue, Fig. 1), uses Efficient (linear) attention [15] for atlas-scale scalability.

**Table 1.** Ablations on the MERFISH mouse primary motor cortex benchmark. Each row changes one component relative to the full G2T model (EDM head, *K*=8 embedding, flow matching, 32 attention heads). Values are mean ± SD over ten seeds (0–9). ^*^/^**^/^***^ mark a difference from the full model significant at *p <* 0.05/0.01/0.001 under a paired *t*-test across the shared seeds; unmarked differences are not significant at *n*=10. Spearman and Contact F1 are higher-is-better; Sum RSSD lower-is-better.

| Variant | Spearman ↑ | Contact F1 ↑ | Sum RSSD ↓ |
| --- | --- | --- | --- |
| **G2T (full)** | **$0.472 \pm 0.005$** | **$0.0624 \pm 0.0008$** | **$75.9 \pm 1.3$** |
| – EDM head (predict 2-D) | $0.460 \pm 0.010^*$ | $0.0612 \pm 0.0009^*$ | $77.6 \pm 1.1^{**}$ |
| Diffusion (vs. flow matching) | $0.469 \pm 0.008$ | $0.0606 \pm 0.0006^{***}$ | $76.5 \pm 1.3$ |
| 16 heads (vs. 32) | $0.464 \pm 0.008^*$ | $0.0611 \pm 0.0007^{**}$ | $76.7 \pm 1.7$ |
| $K=2$ (vs. $K=8$) | $0.455 \pm 0.012^{**}$ | $0.0607 \pm 0.0012^*$ | $78.3 \pm 2.0^*$ |
| $K=16$ (vs. $K=8$) | $0.469 \pm 0.005$ | $0.0621 \pm 0.0006$ | $76.1 \pm 1.2$ |

**Figure 1.**
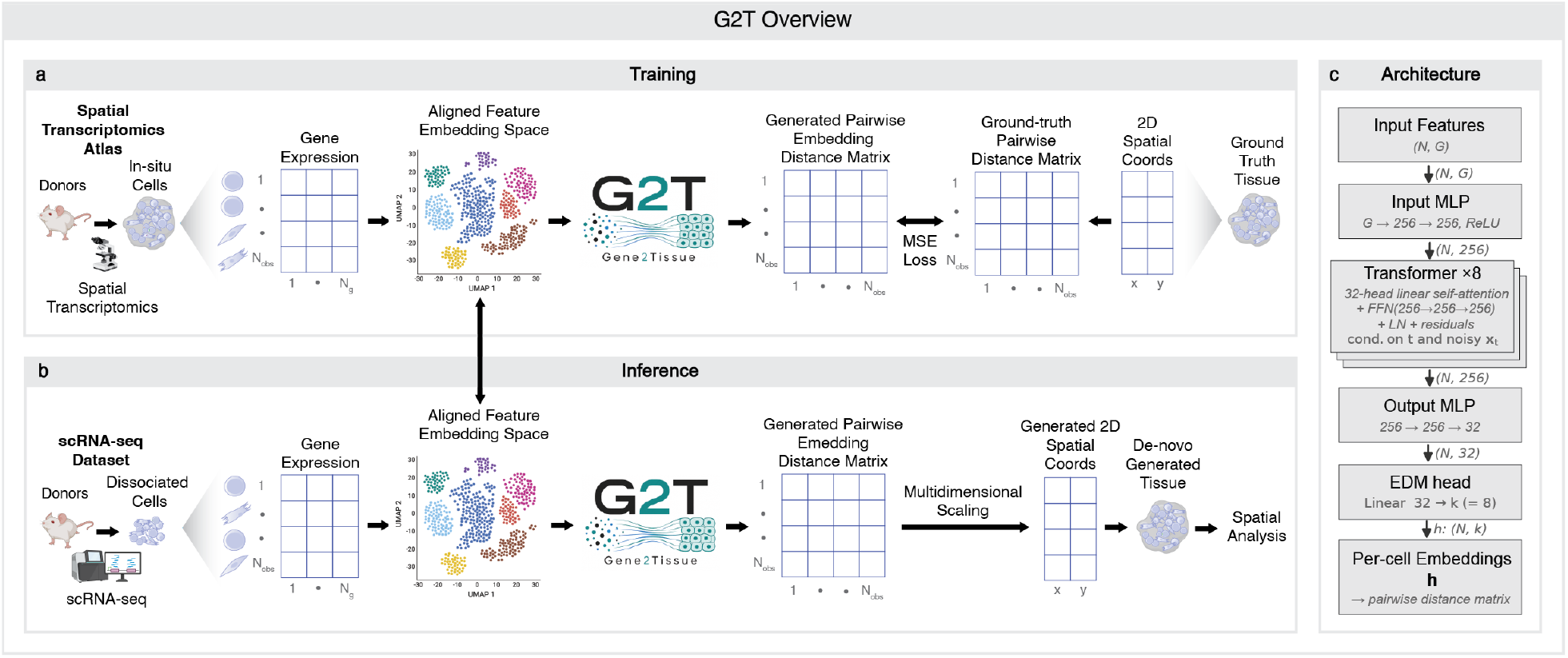
Overview of G2T. **(a)** *Training*. Gene expression is encoded per cell and decoded to a pairwise embedding-distance matrix, matched by MSE to the ground-truth distance matrix derived from the 2-D coordinates. **(b)** *Inference*. Dissociated scRNA-seq cells pass through the same network; Fast MDS (LOBPCG) turns the predicted distance matrix into a 2-D tissue reconstruction. **(c)** *Architecture*. An input MLP (*G* → 256 → 256), eight Transformer blocks with 32-head Efficient (linear) self-attention and FFN (256 → 256 → 256) conditioned on the flow-matching time *t* and noisy positions **x**_*t*_, an output MLP (256 → 256 → 32), and an EDM head (linear 32 → *K*=8) yield per-cell embeddings **h** ∈ ℝ^*N*×*K*^ whose pairwise squared distances define the predicted distance matrix.

**Contributions**. (i) We introduce *embedding-distance flow matching*: flow matching runs on the cell coordinates, but the denoiser is reparameterised — via an Euclidean-distance-matrix (EDM) head to predict per-cell embeddings whose pairwise squared distances define the target geometry rather than regressing 2-D coordinates, relaxing the 2-D output parameterisation of prior diffusion-based methods and yielding a frame-invariant target. (ii) We pair this with a Locally Optimal Block Preconditioned (LOBPCG [7]) MDS read-out that recovers 2-D coordinates from the predicted distance matrix via a top-2 eigendecomposition. (iii) On the MERFISH mouse primary motor cortex benchmark [24], G2T improves over LUNA — the prior state of the art — across all three reported metrics (+4.2% Spearman and +5.0% Contact F1, higher better; −1.6% Sum RSSD, lower better). (iv) On the mouse central nervous system scRNA-seq atlas — evaluated against a STARmap PLUS-imputed spatial reference rather than measured coordinates [16] — G2T again outperforms LUNA and CeLEry, with even larger relative margins (+14.7% Spearman and +7.9% Contact F1, higher better; −6.6% Sum RSSD, lower better).

## 2. Method

### 2.1. Problem formulation

Consider a tissue slice with *N* cells, gene expression **X** ∈ℝ^*N*×*G*^ and 2-D coordinates **Y** ∈ ℝ^*N*×2^. At training time we observe both **X** and **Y**; at inference time on a query slice or scRNA-seq dataset only **X** is available and our goal is to recover the spatial arrangement of cells. Because any rigid motion of **Y** is biologically equivalent [24], we supervise the model on the rigid-motion invariants of **Y**: its *N* ×*N* pairwise squared-distance matrix 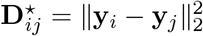.

### 2.2. Architecture

G2T (Fig. 1c) is built from four components.

#### Input projection

Each cell’s expression vector **x**_*i*_ ∈ ℝ^*G*^ is projected by a two-layer MLP *G* → 256 → 256 (ReLU) to an initial latent 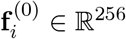.

#### Transformer trunk

Eight Transformer blocks [20] refine the latents using a 32-head Efficient Attention [15] that has linear (rather than quadratic) time and memory complexity in *N*, letting us process the largest atlas sections (tens of thousands of cells) whole. The pairwise-distance loss and MDS read-out are still *O*(*N* ^2^) per slice, so this removes attention’s quadratic cost without making the method linear-scale end-to-end. Each block contains LayerNorm, multi-head linear self-attention, a position-wise FFN (256 → 256 → 256) and residual connections. Following the LUNA architecture, every block is conditioned on the current flow-matching time *t* (via a learned time embedding *γ*_*t*_) and on the current noisy positions **x**_*t*_ (via an auxiliary position stream), allowing the embedding update to depend on how far the trajectory has progressed.

#### Output MLP

A two-layer MLP 256 → 256 → 32 projects the final latent into a lower-dimensional space.

#### EDM head

The head outputs per-cell *embeddings* **h**_*i*_ ∈ ℝ^*K*^ (*K*=8) via a linear layer 32 → *K*; “EDM” names the Euclidean-distance-matrix these embeddings induce (and the loss applied to it), not a direct matrix output. The predicted matrix is then

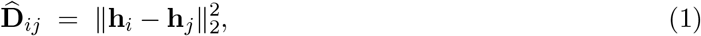

which is differentiable in **h** and, by construction, invariant to any isometry of the embeddings **h** (rotation, translation, or reflection). Predicting a pairwise-distance representation and recovering coordinates from it is the classical route of Euclidean-distance-matrix geometry [2] and mirrors distance-based structure prediction, e.g. AlphaFold [5], which predicts a distogram of inter-residue distances and then recovers 3-D structure from it.

### 2.3. Training objective

We train G2T with conditional flow matching [11], with the probability path defined on the cell *coordinates*. For each training slice we sample *t* ∼ Uniform(*t*_min_, 1) (*t*_min_=10^−3^), draw **ε** ∼ N (**0, I**) and corrupt the coordinates as **x**_*t*_ = (1 − *t*) **y** + *t* **ε** (so **x**_*t*_→**y** as *t*→0; sampling integrates the reverse ODE from *t*=1 to 0). Conditioned on **x**_*t*_, the time *t* and the gene expression **X**, the network produces through the EDM head — per-cell embeddings **h**_*i*_ ∈ ℝ^*K*^ (*K*=8) whose pairwise squared distances form the predicted distance matrix 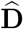 (Eq. 1). G2T uses the data-prediction (x_0_) parameterisation of flow matching: 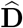 defines an estimate yb0 of the clean coordinates — recovered by the MDS read-out of §2.4 — with induced velocity field **v**_*θ*_(**x**_*t*_, *t*) = (**x**_*t*_ − **ŷ**_0_)*/t*. On the straight-line path, regressing **ŷ**_0_ toward the clean coordinates is equivalent — up to the standard 1*/t*^2^ weighting — to regressing the conditional flow-matching velocity **ε** − **y**. Because a cell cloud is defined only up to a rigid motion, however, we do not penalise the raw-coordinate error ∥**ŷ**_0_ − **y**∥^2^ (which would charge for the arbitrary global frame) but its SE(2)-invariant surrogate, the pairwise squared-distance error between 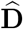 and the ground-truth matrix **D**^*⋆*^:

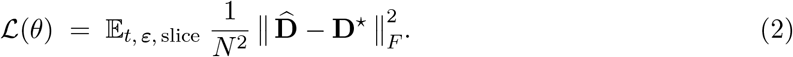

Since 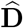 and D⋆ are the isometry invariants of Ŷ0 and y, this surrogate has the same clean target as the **x**_0_-regression objective but quotients out the arbitrary global frame: it vanishes exactly when **ŷ**_0_ equals **y** up to a planar Euclidean isometry — a rotation, translation, *or reflection*, since equal pairwise distances cannot distinguish a mirror image (our Procrustes step allows reflection, so mirrored reconstructions are treated as equivalent). We therefore keep the flow-matching probability path, the **x**_0_ parameterisation, and the rectified-flow sampler (§2.4) unchanged and change only the objective. This invariance is of the *representation* (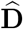is unchanged by isometries of the *K*-dimensional embeddings), leaving the recovered 2-D tissue fixed only up to the biologically relevant planar rigid motion (as in LUNA [24]); the **x**_0_–velocity equivalence above makes this a change of *objective*, not of the generative process. The *K*=8 embedding relaxes the output parameterisation rather than adding intrinsic dimensionality: the ideal minimiser of the distance loss is still a 2-D-embeddable matrix (as **D**^*⋆*^ derives from 2-D coordinates), but predicting an unconstrained *K*-dimensional EDM — instead of being forced to emit an exactly-2-D-embeddable output at every denoising step — is an over-parameterisation that empirically eases optimisation (Table 1: *K*=2 falls to the LUNA level while *K*=8 is best).

### 2.4. Inference: MDS read-out and reverse ODE

We integrate the reverse ODE from *t*=1 (Gaussian noise) to *t*→0 over 50 uniform Euler steps, conditioned on the query gene expression. At each step the predicted matrix 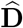 is decoded to a coordinate estimate **ŷ**_0_ by classical MDS [17] on the double-centred Gram matrix 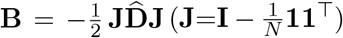 as only the top-2 eigenpairs are needed we use the LOBPCG eigensolver [7] (“Fast MDS”), avoiding the *O* (*N* ^3^) full diagonalisation. MDS determines ŷ0 only up to a rigid motion, so we Procrustes/Kabsch-align [6] it to the current iterate **x**_*t*_ to fix the frame. For numerical stability we add a small Tikhonov diagonal before the top-2 eigendecomposition and, for any slice with fewer than three cells (too few to embed in 2-D), keep its current iterate instead. We form 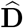 via the memory-efficient expansion 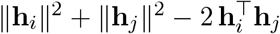 rather than by squaring differences, so a non-negativity clamp absorbs the finite-precision round-off this incurs for near-coincident embeddings. These stability-only safeguards rarely engage, since the distance loss is minimised by a valid, 2-D-embeddable EDM (full details in the Supplementary Material). Intuitively the read-out stays faithful because the target **D**^*⋆*^ is itself the distance matrix of 2-D coordinates: the loss only rewards embeddings whose pairwise distances are realisable in the plane, so the extra *K*=8 dimensions give the denoiser slack but are squeezed out as it converges. Empirically the learned geometry is nearly two-dimensional: across the 31 cortex test sections the top two eigenvalues of **B** capture 99.2% of its positive spectral mass at the final denoising step (SD 0.1%, range 98.8–99.4%), so the top-2 read-out discards under 1% of the structure — confirming that the *K*=8 embedding relaxes the output parameterisation without pushing the recovered geometry off the plane. A single deterministic Euler step of the rectified-flow ODE then advances the coordinates from time *t* to *s < t*,

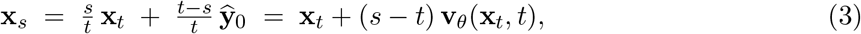

which is exact for a *fixed* predicted endpoint under the rectified-flow path (the velocity is then constant along the trajectory) and returns **ŷ**_0_ at *s*=0. In practice the network re-conditions on **x**_*t*_ and re-estimates ŷ0 at each step, so the reverse process is an Euler integration of a *learned* velocity field — a genuine iterative refinement, not a one-shot distance-matrix regression.

### 2.5. Application to dissociated scRNA-seq

At inference G2T requires only gene expression as observed input; the coordinates it conditions on are initialised from noise and evolved by the reverse process, never supplied. Inference on dissociated scRNA-seq is therefore mechanically identical to inference on a held-out ST slice. When query and training data come from different platforms, we follow LUNA [24] and Harmony-integrate [8] the joint expression matrix on shared genes before encoding. The resulting 2-D layout feeds directly into spatial-analysis tools that expect coordinates [13, 19, 4].

### 2.6. Implementation

G2T is implemented in PyTorch. Expression matrices are loaded from AnnData [21] files, log_2_-normalised following LUNA [24]. The cortex model takes the 254 quality-controlled genes of the 258-gene MERFISH panel [25] as input; the cross-platform CNS model instead takes a shared 600-dimensional Harmony [8] latent over the 804 genes common to the ABC [26] and CNS panels — not the raw genes. Coordinates are min–max normalised per slice to [−0.5, 0.5]. We train with AdamW (lr 5×10^−4^) and an exponential moving average of the weights (decay 0.95); the cortex model is trained for 1,000 epochs and the CNS model for 3,500, matching the LUNA budget [24].

## 3. Experiments

### 3.1. MERFISH mouse primary motor cortex

We follow the LUNA cortex split [24]: 33 sections (158,379 cells, Donor 1) train, 31 sections (118,036 cells, Donor 2) test (Fig. 2a). We report LUNA’s three metrics: per-cell Spearman correlation of pairwise distance ranks aggregated as the mean across slices of per-slice medians; Contact F1 on the closest 1% of cell pairs; and Sum RSSD, the sum over cell classes of each class’s Kabsch [6]-aligned root-sum-of-squared coordinate deviation (lower is better; a degenerate collapse inflates it, so CeLEry sets the Sum-RSSD axis scale in Figs. 2–3, where G2T/LUNA sit near 76; cf. the full-model row of Table 1). Because Sum RSSD is a root-*sum* of per-cell-class alignment residuals in the per-slice-normalised frame (tissue width 1), its absolute scale grows with cell count and is not directly interpretable; we therefore read it comparatively and treat a relative reduction as meaningful only when it is consistent across seeds and echoed by the bounded per-cell metrics — Spearman (rank correlation of pairwise distances) and Contact F1, the most intuitive of the three, for which the +5.0% cortex gain means 5% more of the true closest-1% neighbour pairs are recovered. All methods share the identical split, seeds, and metric implementation, with LUNA and CeLEry run from their released code. Each method is trained for five random seeds (0–4); bars report the mean across seeds. LUNA’s benchmark covers novoSpaRc [12], Tangram [1], CytoSPACE [18], STEM [3], scSpace [14], STALocator [9] and CeLEry [27], of which LUNA is by far the strongest; we re-run LUNA and CeLEry here and take the other methods’ standings from that published benchmark rather than re-running them (so those numbers inherit its preprocessing and metric choices). We focus the head-to-head on LUNA and CeLEry (Fig. 2b).

**Figure 2.**
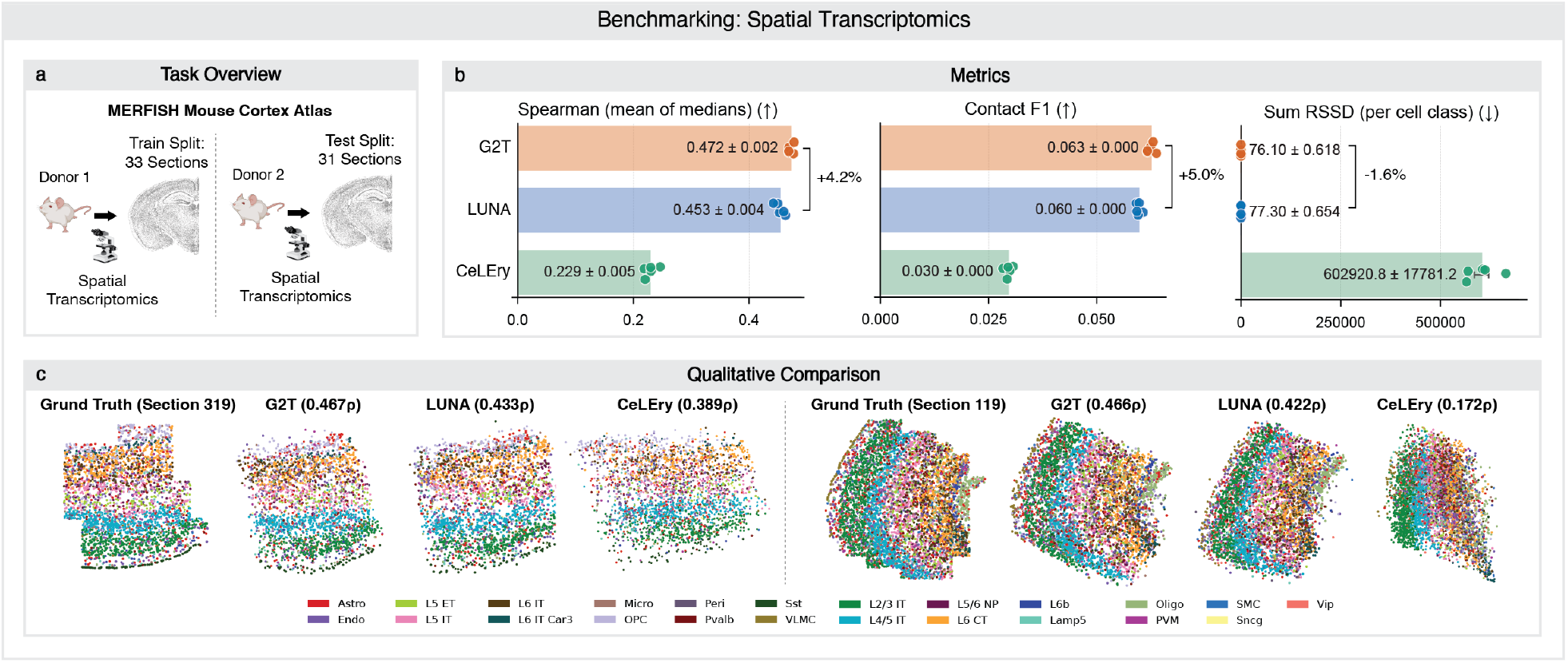
MERFISH mouse primary motor cortex benchmark [25]. **(a)** Split following LUNA [24]: 33 Donor-1 sections train, 31 held-out Donor-2 sections test. **(b)** G2T vs. LUNA [24] and CeLEry [27] on Spearman (mean of per-slice medians), Contact F1 (closest 1% of pairs), and per-cell-class Sum RSSD after Kabsch alignment [6]; G2T improves over LUNA by +4.2% and +5.0% on Spearman and Contact F1 (higher is better) and by −1.6% on Sum RSSD (lower is better). Bars: mean ± SEM over five seeds (0–4). **(c)** Reconstructions of two held-out sections; colours denote cortical cell classes, parentheses give per-section Spearman.

G2T improves over LUNA on all three metrics, raising Spearman from 0.453 to 0.472 (+4.2%), Contact F1 from 0.060 to 0.063 (+5.0%), and reducing per-cell-class Sum RSSD by 1.6%; CeLEry trails both by a large margin. Contact F1 is evaluated on only the closest 1% of cell pairs — a stringent local-contact criterion — so absolute values near 0.06 are expected and lie in LUNA’s reported regime; G2T is the best of the compared methods, though whether this local-contact fidelity suffices for a given downstream niche or communication analysis remains to be established. Qualitative reconstructions of two representative held-out sections (Fig. 2c) show G2T recovers the layered cortical architecture and cell-class compartmentalisation more faithfully than either baseline, with CeLEry collapsing tissue into a small region.

### 3.2. Mouse central nervous system scRNA-seq atlas

We next evaluate the scRNA-seq transfer scenario of LUNA [24], the closest realistic setting to typical dissociated atlases with no measured spatial information. We train G2T on the MERFISH ABC mouse-brain atlas (Animal 1, 147 sections, 2.85 M cells) [26] and apply it to the mouse CNS scRNA-seq atlas [16] (1.08 M cells, 14 held-out reference sections). To bridge the two gene panels we restrict to their 804 common genes and Harmony-integrate [8, 23] them into a shared 600-dimensional latent that serves as the model input, following LUNA. Because the CNS atlas lacks measured spatial coordinates, we follow LUNA’s protocol and evaluate against an *imputed* spatial reference — the locations Shi et al. [16] assigned to these cells by STARmap PLUS integration — rather than measured ground truth; G2T never sees this reference. This scores agreement with a spatial-integration procedure rather than observed coordinates, so the CNS numbers are best read as a relative method comparison, not absolute accuracy. Across five seeds, G2T improves over both LUNA and CeLEry on all three metrics (Fig. 3b), raising Spearman from 0.157 to 0.180 (+14.7%) and Contact F1 from 0.038 to 0.041 (+7.9%), and reducing per-cell-class Sum RSSD by 6.6%; the relative gains over LUNA are markedly larger than on the cortex benchmark. Qualitative reconstructions of two held-out sections (Fig. 3c) show G2T recovering the coarse tissue organisation more faithfully than either baseline — with a higher per-section Spearman than both in each example — while CeLEry degenerates to a near-random layout.

**Figure 3.**
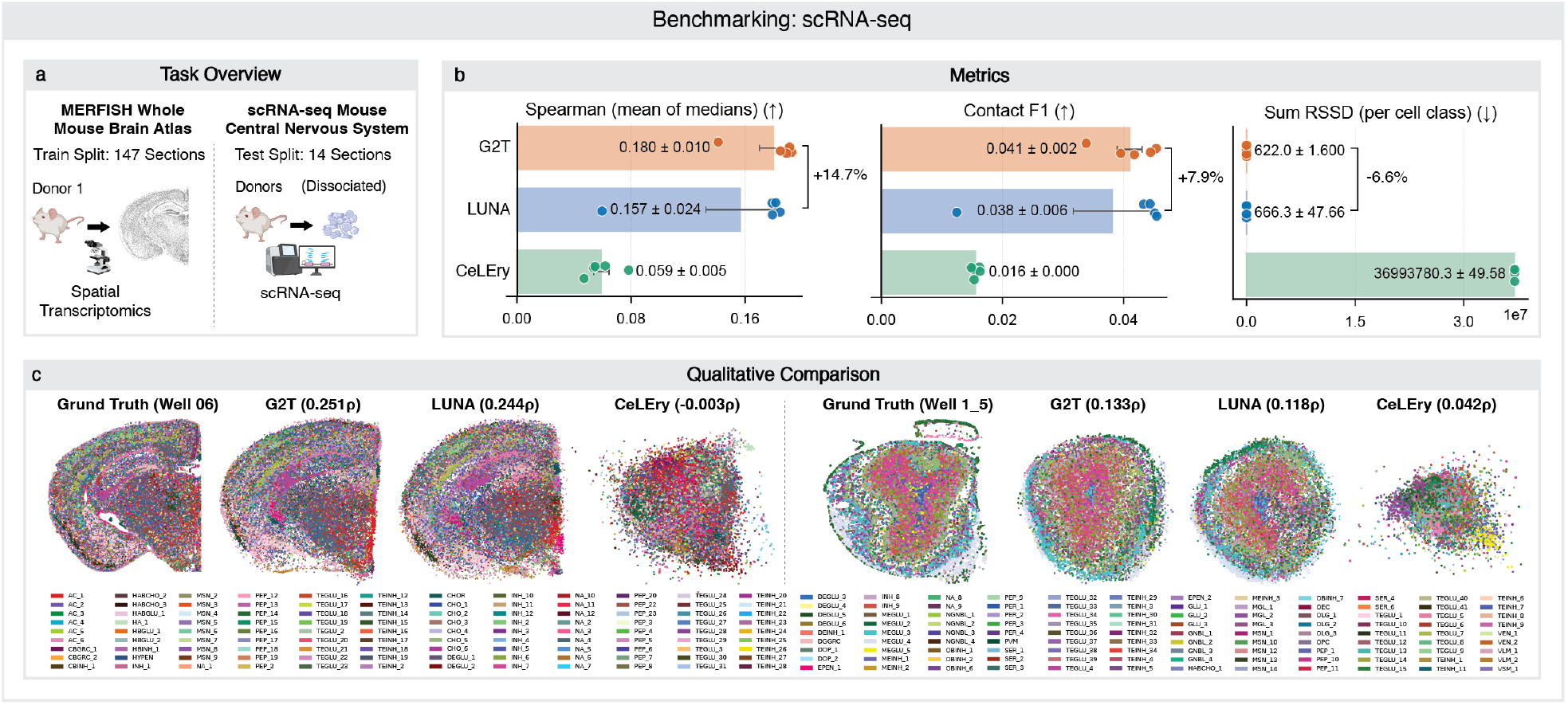
Mouse central nervous system scRNA-seq benchmark. **(a)** Trained on the MERFISH ABC mouse-brain atlas (Animal 1, 147 sections) [26], applied to the CNS scRNA-seq atlas (1.08 M cells, 14 test sections), Harmony-integrated [8] on 804 common genes into a 600-dim latent; the evaluation target is an *imputed* spatial reference (STARmap PLUS-integrated locations [16]), *not* measured coordinates. **(b)** G2T vs. LUNA [24] and CeLEry [27]; G2T improves over LUNA by +14.7% and +7.9% on Spearman and Contact F1 (higher is better) and by −6.6% on Sum RSSD (lower is better; CeLEry’s value sets the axis scale). Bars: mean ± SEM over five seeds (0–4). **(c)** Reconstructions of two held-out sections (the reference panels show the *imputed* STARmap PLUS locations, not measured ground truth); parentheses give per-section Spearman.

### 3.3. Ablations

We ablate G2T’s main components on the cortex benchmark, changing one factor at a time and re-training ten seeds (0–9) each (Table 1); the full-model row closely matches the headline cortex result.^1^ Every ablation is worse than, or on par with, the full model, and none improves it. Because all variants share the same ten seeds, we compare each to the full model with a paired *t*-test. Removing the EDM head — regressing 2-D coordinates directly, as in prior coordinate-denoising diffusion models — significantly worsens all three metrics (Spearman and Contact F1 *p <* 0.05, Sum RSSD *p*=0.004), directly validating the distance-matrix reparameterisation as G2T’s central design choice. The embedding-width sweep tells the same story: collapsing the embedding to *K*=2 (the 2-D output width) is significantly worse than *K*=8 on all three metrics (Spearman *p*=0.005), whereas widening to *K*=16 is statistically indistinguishable from *K*=8 (all *p >* 0.18) — so lifting the target just above the 2-D bottleneck captures the gain, and *K*=8 is a compact operating point on the resulting plateau. Halving the attention heads to 16 significantly lowers Spearman (*p*=0.01) and Contact F1 (*p*=0.002). Replacing flow matching with a diffusion model at the standard 1,000-step DDPM schedule (LUNA’s setting) significantly lowers Contact F1 (*p <* 0.001) but matches the full model on Spearman and Sum RSSD — so flow matching loses no accuracy while needing far fewer sampling steps (50 Euler steps vs. 1,000).

## 4. Conclusion

We presented G2T, a generative model for tissue reassembly: flow matching on cell coordinates with the denoiser reparameterised through an embedding-distance (EDM) head that lifts the target from 2-D coordinates into a higher-dimensional (*K*=8) space, so the supervised objective is the frame-invariant pairwise-distance matrix and coordinates are recovered by LOBPCG-based MDS. On the MERFISH mouse cortex benchmark and the mouse CNS scRNA-seq atlas, G2T improves over LUNA and CeLEry across all three reported metrics. A promising direction is to better align scRNA-seq and ST expression measurements (e.g. via stronger batch correction or domain adaptation), closing the train/inference input gap. More broadly, we expect embedding-distance flow matching to be a useful primitive for spatial-omics learning.

## Footnotes

1 The headline cortex numbers (Fig. 2) average five seeds (0–4), whereas the ablations here average ten (0–9); the full-model ablation row therefore differs slightly from the headline (Contact F1 0.0624 vs. 0.063; Sum RSSD 75.9 vs. 76.1). Spearman coincides at 0.472.

