## Supplementary Material for "G2T: Tissue Reconstruction from Gene Expression via Embedding-Distance Flow Matching"

Proceedings of Machine Learning Research 1:1–2, 2026 Machine Learning in Computational Biology (MLCB) 2026

### 1. Implementation and reproducibility details

**Hardware and code.** All models are trained on a single NVIDIA H200 GPU. Source code is available at `https://github.com/Lotfollahi-lab/g2t`, and code to reproduce the experiments at `https://github.com/Lotfollahi-lab/g2t-reproducibility`.

**Data and batching.** Training uses *whole* tissue slices with no cell subsampling; a batch is 6 slices (cortex slices hold $\sim$7k cells each; the ABC/CNS slices are larger). Expression is $\log_2(1+x)$-normalised and cell coordinates are min–max normalised per slice to $[-0.5, 0.5]$. Cross-platform runs use the *intersection* of the two gene panels (the 804 genes common to the ABC MERFISH and CNS panels; no missing-gene imputation), Harmony-integrated [2] into a shared 600-dimensional latent that replaces the raw expression as the model input.

**Architecture and conditioning.** The backbone is LUNA's three-stream transformer (node-feature / time / position streams): an input MLP $G \to 256 \to 256$, eight blocks with 32-head Efficient (linear) attention [3] (LayerNorm, position-wise FFN $256 \to 256 \to 256$, residual connections), an output MLP $256 \to 256 \to 32$, and the linear EDM head $32 \to K{=}8$. Each block is conditioned on the flow-matching time $t$ via a learned embedding $\gamma_t$ and on the current noisy positions $\mathbf{x}_t$ via the auxiliary position stream.

**Training and inference.** We use AdamW (lr $5 \times 10^{-4}$), an exponential moving average of the weights (decay 0.95), and the $\mathbf{x}_0$-prediction linear (rectified-flow) path with $t \sim \mathcal{U}(10^{-3}, 1)$; the cortex model trains for 1,000 epochs and the CNS model for 3,500. Inference integrates the reverse ODE for 50 Euler steps and recovers coordinates by MDS *at every step* (not only at the end): the predicted squared-distance matrix is double-centred, its top-2 eigenpairs are extracted by LOBPCG (default) or exact `eigh`, and the coordinates are Procrustes/Kabsch-aligned [1] to $\mathbf{x}_t$. Numerical guards: because the pairwise squared distances are formed via the identity $\|\mathbf{h}_i - \mathbf{h}_j\|^2 = \|\mathbf{h}_i\|^2 + \|\mathbf{h}_j\|^2 - 2\,\mathbf{h}_i^\top \mathbf{h}_j$ (which avoids an $N \times N \times K$ intermediate at atlas scale), we clamp them to be non-negative to remove the tiny negatives finite precision produces for near-coincident embeddings; a small Tikhonov term stabilises the eigendecomposition; and any slice with fewer than three cells (or a degenerate Gram matrix) keeps its current-iterate layout instead of the MDS output. To quantify how well the learned distance matrix admits a 2-D embedding, we log at every reverse step the fraction $(\lambda_1 + \lambda_2)/\sum_{\lambda>0} \lambda$ of the double-centred Gram matrix's positive spectral mass carried by its top two eigenvalues (a full `eigh`, independent of the deployed solver). At the final denoising step this fraction is 99.2% on average across the 31 cortex test sections (SD 0.1%, range 98.8–99.4%), so the top-2 read-out is near-lossless despite the $K{=}8$ embedding width.

© 2026

**Metrics.** Per-cell Spearman is the correlation of predicted and true pairwise-distance ranks, aggregated as the mean over slices of per-slice medians; Contact F1 is the F1 over the closest 1% of cell pairs; and Sum RSSD sums, over cell classes, each class's Kabsch-aligned root-sum-of-squared coordinate deviation,

$$\text{Sum RSSD} = \sum_c \sqrt{\textstyle\sum_{i \in c} \|\mathbf{R}_c \,\widehat{\mathbf{y}}_i - \mathbf{y}_i\|_2^2},$$

where $\mathbf{R}_c$ is the per-class optimal rotation (lower is better). This per-class alignment is a property of the *metric*, not of inference: a cell class with fewer than two cells in a slice — too few to fit a rotation — is excluded from that slice's Sum RSSD (this is distinct from the per-slice MDS fallback above).

**Baselines.** LUNA and CeLEry are run from their released code with recommended settings, on the identical split, seeds, and metric implementation.

[3] Zhuoran Shen, Mingyuan Zhang, Haiyu Zhao, Shuai Yi, and Hongsheng Li. Efficient attention: Attention with linear complexities. In *IEEE/CVF Winter Conference on Applications of Computer Vision (WACV)*, 2021.
